# Homologous and heterologous serological response to the N-terminal domain of SARS-CoV-2

**DOI:** 10.1101/2021.02.17.431722

**Authors:** Huibin Lv, Owen Tak-Yin Tsang, Ray T. Y. So, Yiquan Wang, Meng Yuan, Hejun Liu, Garrick K. Yip, Qi Wen Teo, Yihan Lin, Weiwen Liang, Jinlin Wang, Wilson W. Ng, Ian A. Wilson, J. S. Malik Peiris, Nicholas C. Wu, Chris K. P. Mok

## Abstract

The increasing numbers of infected cases of coronavirus disease 2019 (COVID-19) caused by severe acute respiratory syndrome coronavirus 2 (SARS-CoV-2) poses serious threats to public health and the global economy. Most SARS-CoV-2 neutralizing antibodies target the receptor binding domain (RBD) and some the N-terminal domain (NTD) of the spike protein, which is the major antigen of SARS-CoV-2. While the antibody response to RBD has been extensively characterized, the antigenicity and immunogenicity of the NTD protein are less well studied. Using 227 plasma samples from COVID-19 patients, we showed that SARS-CoV-2 NTD-specific antibodies could be induced during infection. As compared to the serological response to SARS-CoV-2 RBD, the SARS-CoV-2 NTD response is less cross-reactive with SARS-CoV. Furthermore, neutralizing antibodies are rarely elicited in a mice model when NTD is used as an immunogen. We subsequently demonstrate that NTD has an altered antigenicity when expressed alone. Overall, our results suggest that while NTD offers an alternative strategy for serology testing, it may not be suitable as an immunogen for vaccine development.

## Introduction

The novel coronavirus SARS-CoV-2, which is the pathogen that has caused the COVID-19 pandemic, has spread to over 216 countries (Liu et al., 2020c). COVID-19 patients show varying disease severity ranging from asymptomatic to requiring intensive care (Liu et al., 2020d). Many studies have now shown that SARS-CoV-2-specific immunoglobulin G (IgG) in COVID-19 patients is a key signature of immune response upon the infection (Barnes et al., 2020; Brouwer et al., 2020; Isho et al., 2020; Jiang et al., 2020; Long et al., 2020; Pinto et al., 2020; Wang et al., 2020). The spike glycoprotein is the immunodominant target for the neutralizing antibody response in COVID-19 patients. Importantly, neutralizing antibodies to the spike are able to maintain detectable levels through at least 5-8 months post-infection (Dan et al., 2021; Lau et al., 2021; Ripperger et al., 2020; Roltgen et al., 2020; Wajnberg et al., 2020). The spike protein consists of S1 (head) and S2 (stem) subunits that are initially connected by a furin cleavage site (Walls et al., 2020). The S1 contains two structurally well-defined domains, namely the N-terminal domain (NTD) and receptor binding domain (RBD). SARS-CoV-2 initiates viral entry by engaging the host receptor angiotensin converting enzyme 2 (ACE2) through the RBD. Most known SARS-CoV-2 neutralizing antibodies to date are RBD-specific (Barnes et al., 2020; Brouwer et al., 2020; Cao et al., 2020; Ju et al., 2020; Liu et al., 2020a; Liu et al., 2020b; Pinto et al., 2020; Rogers et al., 2020; Seydoux et al., 2020; Shi et al., 2020; Wu et al., 2020; Zost et al., 2020). Thus, detection of RBD-specific antibodies is widely used in many serodiagnosis tests (Perera et al., 2020; Premkumar et al., 2020). RBD has also been a major focus in vaccine design (Dai et al., 2020; Tai et al., 2020; Zang et al., 2020). In contrast, the immunogenicity and antigenicity of other domains on the spike is not very well characterized. An increasing number of neutralizing antibodies to the NTD have recently been identified from COVID-19 patients (Cerutti et al., 2021; Chi et al., 2020; Liu et al., 2020b; McCallum et al., 2021; Noy-Porat et al., 2021; Suryadevara et al., 2021; Wang et al., 2021a). In addition, tyrosine-protein kinase receptor UFO (AXL) is suggested to be a co-receptor for SARS-CoV-2 by interacting with the NTD (Wang et al., 2021b). Another recent finding shows that the NTD can interact with tetrapyrrole products that reduce the reactivity of the SARS-CoV-2 spike with human immune sera as a possible mechanism to evade antibody immunity (Rosa et al., 2021). It is thus believed that neutralizing antibodies to NTD antibodies may play an important role in protection against SARS-CoV-2. However, the NTD-specific antibodies have been mainly identified from clonal B cells of individuals. The serological response to the NTD in COVID-19 patients, as well as the immunological properties of NTD are not yet well understood. In this study, we evaluated the human serological response to NTD protein from 227 specimens collected from 141 COVID-19 patients. The cross-reactivity of NTD-specific antibody response to different coronaviruses was also examined. We also explored the serological response by using NTD as an immunogen for immunization in mice.

## Results

### Human serological responses to the NTD of SARS-CoV-2

We tested 227 plasma samples from 141 RT-PCR confirmed COVID-19 patients in Hong Kong and another 195 plasma samples from healthy blood donors that were collected prior to the emergence of SARS-CoV-2 as baseline controls. The samples were tested in parallel in ELISA assays for the IgG against NTD and RBD. For each assay, samples were defined as seropositive if the detection signal was three standard deviations above the mean of baseline controls. There was a progressive increase of seropositivity in the NTD ELISA after the first day of symptom onset, with 25% (12 out of 48) being positive in the first two weeks and 84.9% (152 out of 179) after day 14 to day 141 (Table 1; Figure 1A). Consistent with our previous study (Perera et al., 2020), the positivity in the RBD assay also progressively increased with time after illness onset, with 58.3% (28 out of 48) specimens positive in the first two weeks of illness onset and 98.3% (176 of 179) after day 14 to day 150?? (Table 1, Figure 1B). Specimens that were found to be positive in the NTD ELISA (n=164) were also positive in the RBD ELISA. In fact, there was a strong correlation between the serological response to NTD and RBD proteins after day 8 of symptom onset (Pearson correlation = 0.78) (Figure 1C).

**TABLE 1:** RBD and NTD ELISA results from the plasma of COVID-19 patients

| TABLE 1: RBD and NTD ELISA results from the plasma of COVID-19 patients |  |  |  |
| --- | --- | --- | --- |
|  | Numbers of sample | RBD positive (%) | NTD positive (%) |
| Days 1-7 | 21 | 12 (57.1) | 1 (4.8) |
| Days 8-14 | 27 | 16 (59.3) | 11 (40.7) |
| Days 15-21 | 12 | 12 (100) | 7 (58.3) |
| Days 22-28 | 14 | 14 (100) | 10 (71.4) |
| Days >28 | 153 | 150 (98.0) | 135 (88.2) |
| Total | 227 | 204 (89.9) | 164 (72.2) |

**Figure 1.**
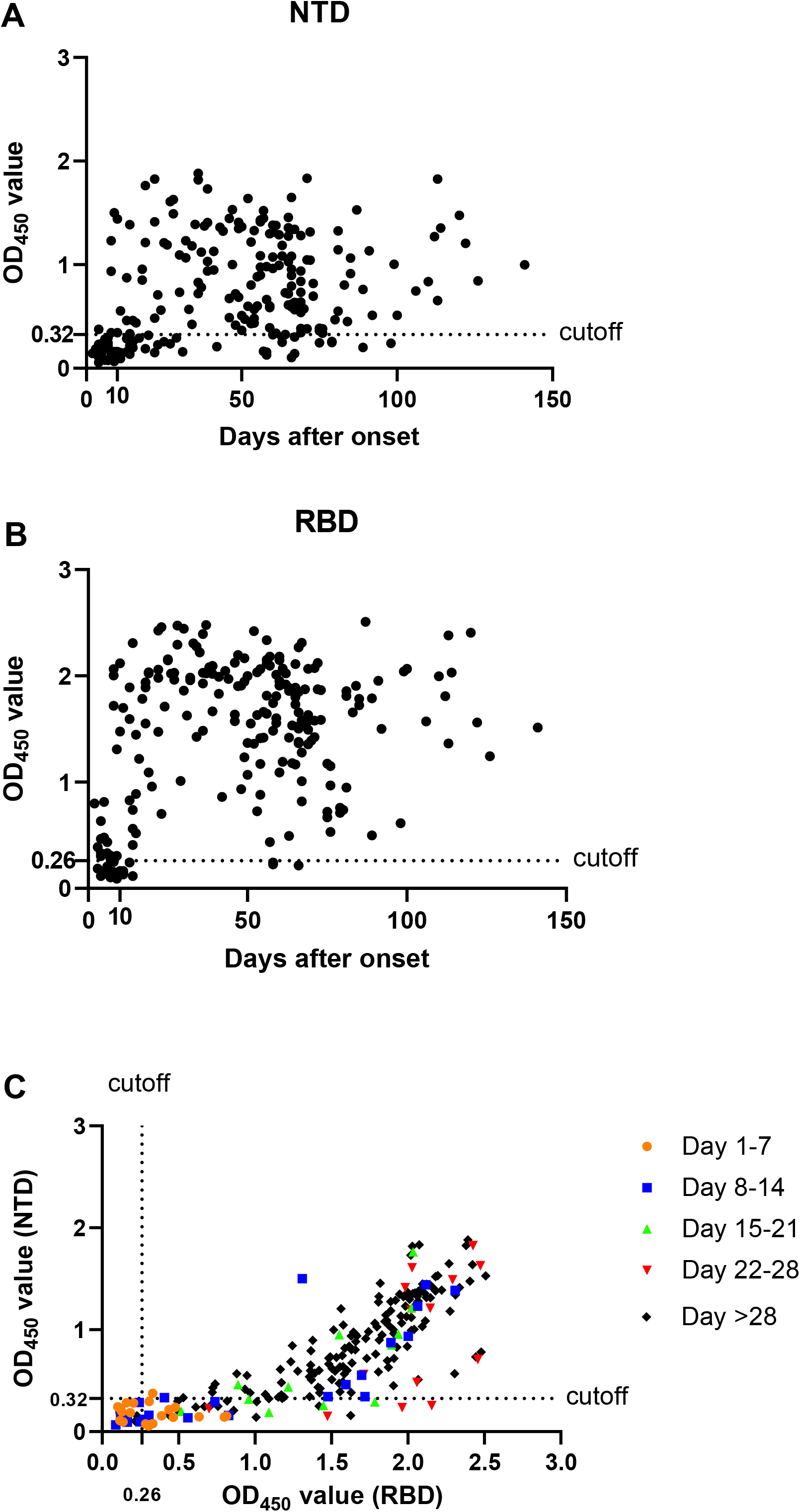
Patient serological responses to SARS-CoV-2 NTD and RBD protein. (A-B) Binding of plasma from SARS-CoV-2 infected patients to SARS-CoV-2 NTD protein (A) and RBD protein (B) were measured during the days symptom after onset by ELISA assay. The mean OD_450_ ELISA binding values calculated after testing each plasma sample in duplicate are shown. The plamsa sample from healthy donors were used as negative control. The ELISA cutoff value of NTD and RBD protein were 0.3272 and 0.2607, respectively (mean + three standard deviations). (C) Pearson correlation (r) was used to assess the relationship between measured SARS-CoV-2 serological binding responses to SARS-CoV-2 RBD and NTD protein in the SARS-CoV-2 infected patients at consequent time periods.

### Cross reactivity of the humoral immunity from COVID-19 patients

The extent of cross-reactive serological responses to other coronaviruses during SARS-CoV-2 infection is not fully understood. Our previous study observed that plasma samples from COVID-19 patients can cross-react with the RBD of SARS-CoV (Lv et al., 2020b). Here, we further tested the binding of 227 plasma samples of COVID-19 patients to the NTDs of SARS-CoV-2 and SARS-CoV. Among the 164 samples with positive binding to the NTD of SARS-CoV-2, only 8 (4.9%) cross-reacted with the NTD of SARS-CoV in the ELISA binding assay (Figure 2A). There is no significant correlation in binding between the groups (Pearson correlation = 0.06). In contrast, among 204 samples that showed positive binding to the RBD of SARS-CoV-2, 158 (77.5%) cross-reacted to the RBD of SARS-CoV. There is a significant correlation in binding between these two RBD antigens (Pearson correlation = 0.43) (Figure 2B). This result is consistent with the RBD having a higher sequence conservation compared to NTD. While the RBDs of SARS-CoV-2 and SARS-CoV share 73% amino-acid sequence identity, their NTDs only share 53% amino-acid sequence identity (Figure S1A-B).

**Figure 2.**
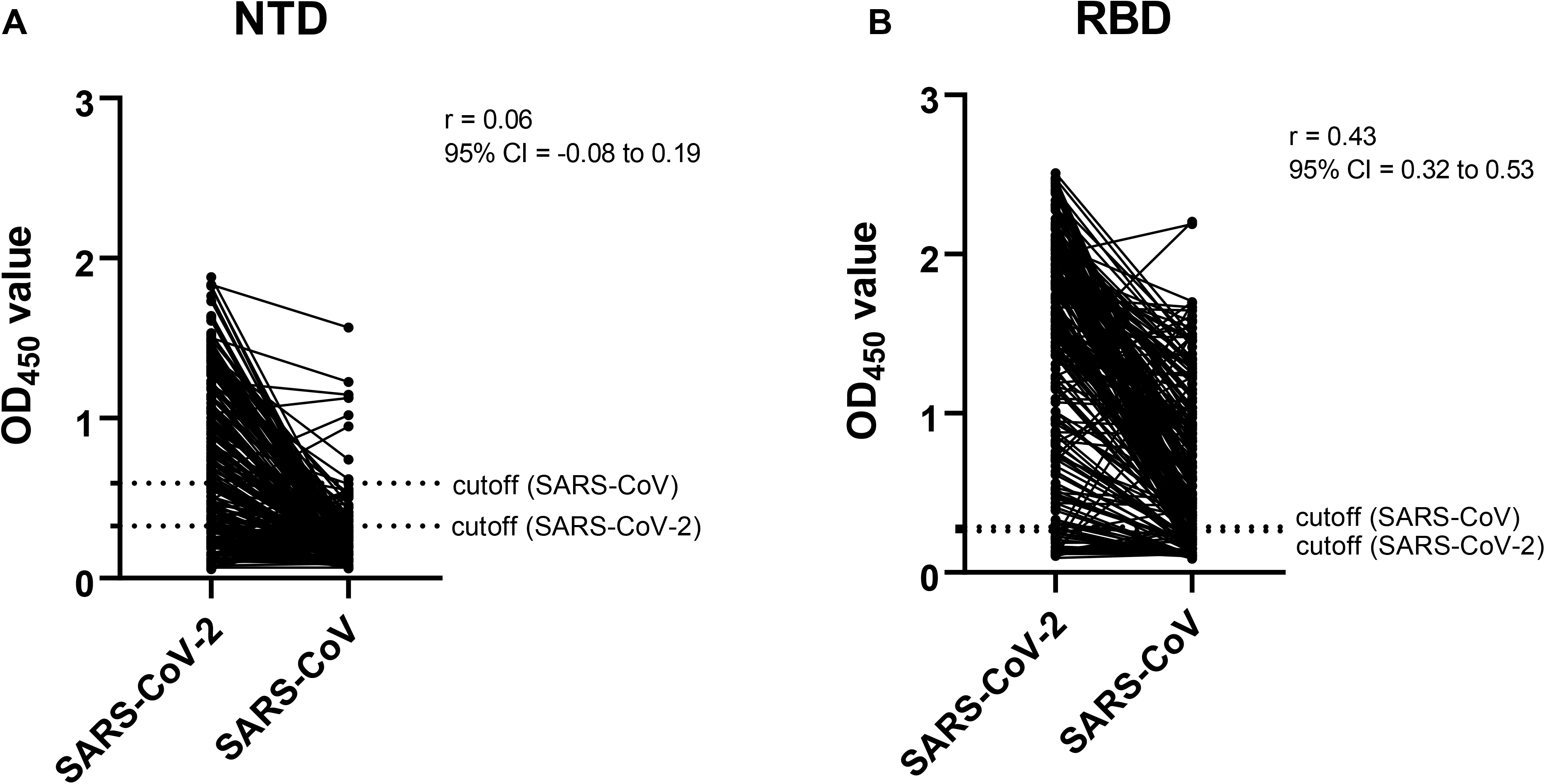
Cross-reactiive serological response to NTD and RBD protein between SARS-CoV and SARS-CoV-2. (A-B) Pearson correlation (r) was used to evaluate the binding capacity of plasma to SARS-CoV and SARS-CoV-2 NTD (A) and RBD (B) protein from 227 SARS-CoV-2 infected patients. The ELISA cutoff value of NTD protein to SARS-CoV and SARS-CoV-2 were 0.5939 and 0.3272, and RBD protein to SARS-CoV and SARS-CoV-2 were 0.2867 and 0.2607, respectively (mean + three standard deviations).

To explore whether SARS-CoV-2 infection can lead to serological responses that cross-react with other human coronaviruses, we selected 118 plasma samples from the COVID-19 patients and tested their binding to the spike proteins of all four known human seasonal coronaviruses, namely 229E, NL63, HKU-1 and OC43. The results were compared to another 118 plasma samples from healthy blood donors that are age- and sex-matched to the COVID-19 cohort. As our control, the plasma of COVID-19 patients showed a significantly higher level of binding to the NTD and RBD of SARS-CoV-2 compared to that of the healthy controls (Figure 3A and B). Compared to the plasma of healthy controls, the plasma of the COVID-19 cohort exhibited significantly higher binding to the spike proteins of HKU1 and OC43 (Figure 3E-F). In contrast, plasma of healthy controls and COVID-19 cohort had only very small differences in binding to the spike proteins of NL63 and 229E, although such a difference for NL63 is still significant (P = 7e-5, two-tailed paired t-test, Figure 3C-D). We also collected longitudinal plasma samples from six COVID-19 patients and tested their binding to the NTD and RBD of SARS-CoV-2 as well as to the spikes of other human coronaviruses by ELISA (Figure S2A-F). Although the increases in binding to the NTD and RBD of SARS-CoV-2 were more dramatic, some patients showed modest elevation of serological responses against the spike of different human coronaviruses, especially HKU1 and OC43. Of note, SARS-CoV, SARS-CoV-2, HKU1 and OC43 are beta-coronavirus, whereas NL63 and 229E are alpha-coronavirus. Our results suggest that memory B cells with epitopes that are conserved among different beta-coronaviruses were boosted after SARS-CoV-2 infection. Consistently, recent studies have shown that antibodies targeting the S2 domain can acquire broad reactivity among beta-coronaviruses (Huang et al., 2021; Sauer et al., 2021).

**Figure 3.**
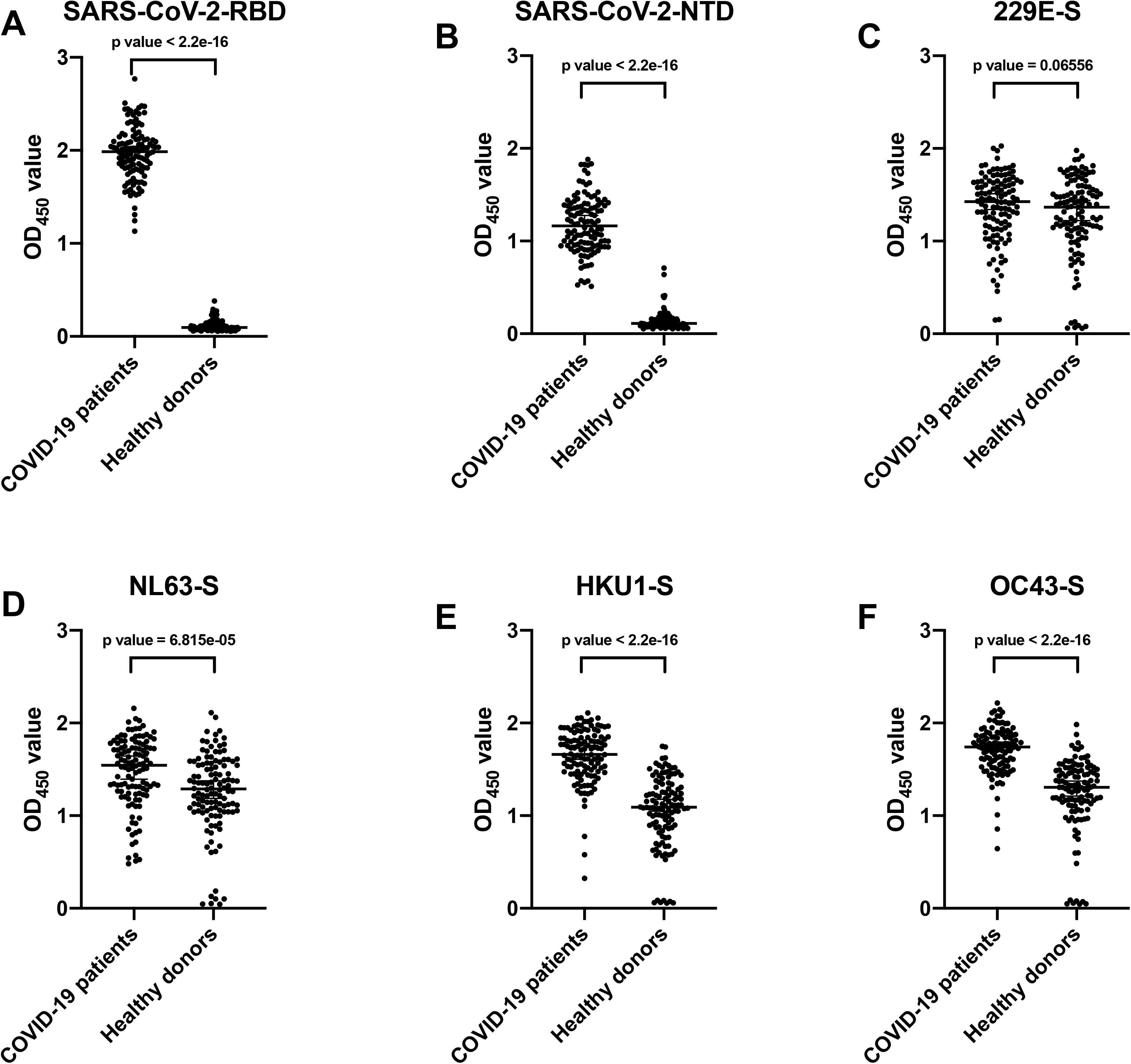
Cross-reactive serological response to human coronaviruses between COVID-19 patients and healthy donors. (A-B) Binding of plasma samples to SARS-CoV-2 NTD (A), SARS-CoV-2 RBD (B), 229E-Spike (C), NL63-Spike (D), HKU1-Spike (E) and OC43-Spike protein (F) were tested by ELISA assay from 118 COVID-2019 patients and age- and sex-matched healthy donors. The OD_450_ value from each dot in the figure was taken by means of two replicates in the same experiment. P-values were caluated using two-tailed paired t-test (***P<0.001). Error bars repeesent strandard deviation.

### Immunization of NTD alone in mice does not induce neutralizing antibody

Since NTD neutralizing antibodies have been shown to confer protection to SARS-CoV-2 (Cerutti et al., 2021; Chi et al., 2020; Liu et al., 2020b; McCallum et al., 2021; Noy-Porat et al., 2021; Suryadevara et al., 2021), we are interested in evaluating if the NTD protein itself is immunogenic and can potentially be a vaccine candidate. We adopted our previous immunization protocol where BALB/c mice were intraperitoneally (i.p.) immunized twice by SARS-CoV-2 or SARS-CoV NTD protein with Addavax as adjuvant (Wu et al., 2019). Plasma samples were collected 14 days after the second immunization and their binding to NTD of SARS-CoV-2 and SARS-CoV was measured by ELISA. We found that immunization with SARS-CoV-2 NTD could induce homologous and cross-reactive binding antibodies to the NTD proteins of SARS-CoV-2 and SARS-CoV (Figure 4A). However, no cross-reactive binding was observed to the SARS-CoV spike protein (Figure 4C). Similarly, plasma samples from mice immunized with SARS-CoV NTD (Figure 4A and C) could cross-react with SARS-CoV-2 NTD protein, but not with the SARS-CoV-2 spike. Although spike binding antibodies could be induced, no viral neutralizing ability could be found after either SARS-CoV or SARS-CoV-2 NTD protein immunization (Figure 4B). As a control, we also tested plasma samples of the mice immunized with live SARS-CoV-2 or SARS-CoV for binding to NTD proteins (Lv et al., 2020a). In contrast to NTD immunization, mice immunized with the live SARS-CoV-2 or SARS-CoV can only elicit NTD antibodies to the autologus strain (Figure 4A and C). No cross-reactivity was found to the spike proteins of NL63, 229E, HKU-1 and OC43 (Figure 4D). These observations suggest that there is a difference in antigenicity between NTD alone and NTD on the spike protein.

**Figure 4.**
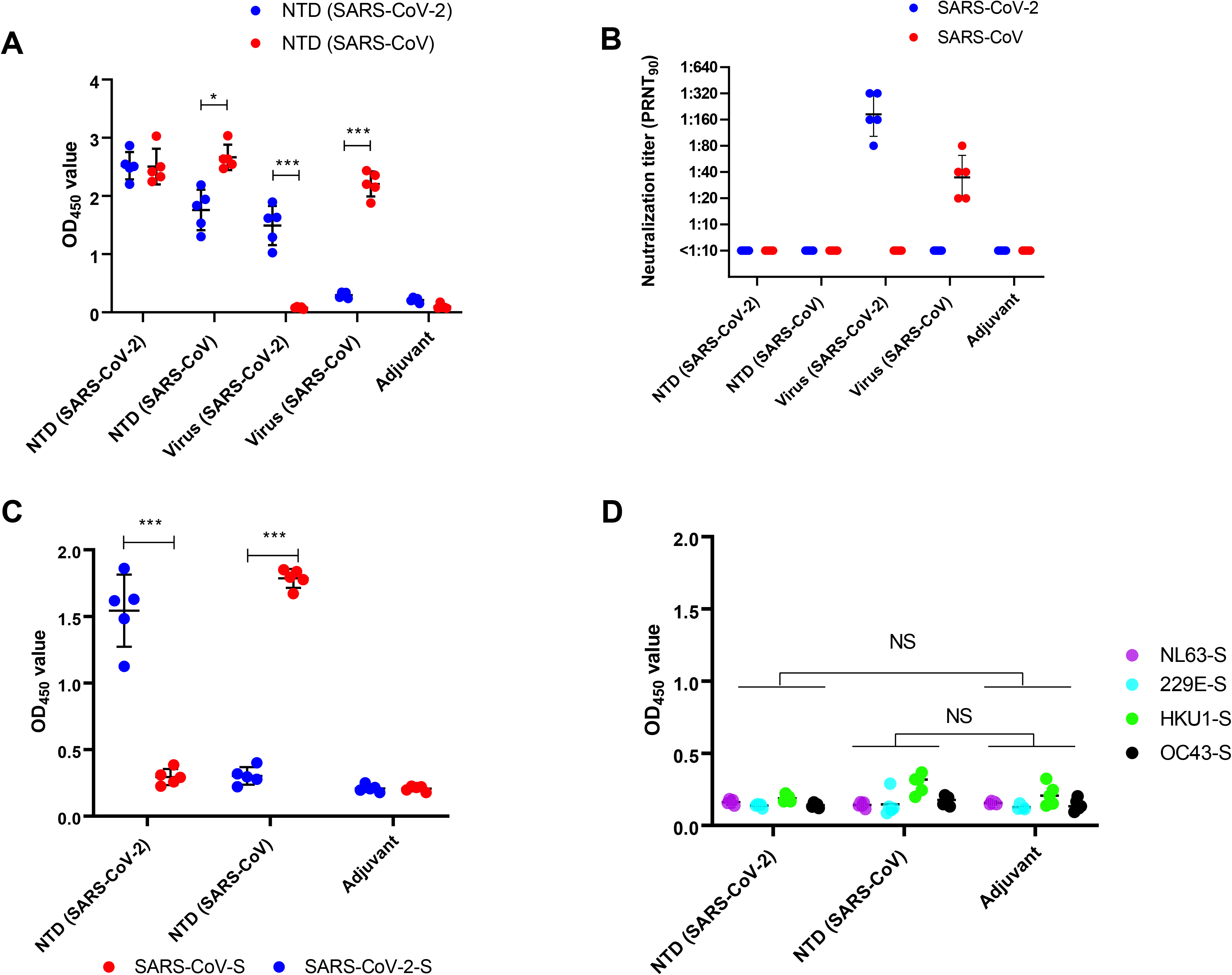
Serological binding and neutralizing capacity against SARS-CoV and SARS-CoV-2 by NTD protein immunization. **(A)** Binding of plasma from SARS-CoV-2 NTD protein immunized mice, SARS-CoV NTD protein immunized mice, live SARS-CoV-2 immunized mice and live SARS-CoV immunized mice against SARS-CoV and SARS-CoV-2 NTD protein were measured by ELISA assay. The mean OD_450_ values calculated after detecting each plasma sample in duplicate are shown. **(B)** Neutralization activities of plasma from mice immunized with SARS-CoV-2 NTD protein, SARS-CoV NTD protein, live SARS-CoV-2 and live SARS-CoV were measured. The value from each dot in the figure was tested by the means of two replicates in the same assay. **(C)** Binding of plasma from SARS-CoV and SARS-CoV-2 NTD protein immunized mice against the full spike of SARS-CoV-2 or SARS-CoV. **(D)** Binding of plasma from SARS-CoV and SARS-CoV-2 NTD protein immunized mice against NL63-Spike, 229E-Spike, HKU1-Spike and OC43-Spike protein were tested by ELISA assay. The OD_450_ value from each dot in the figure was taken by means of two replicates in the same experiment. P-values were caluated using two-tailed t-test. Error bars represent standard deviation.

### A putative structural mechanism of altered antigenicity in NTD alone

To further understand the mechanism of differential antibody responses between immunizations with NTD alone and live virus, we performed a structural analysis of the NTD. A cluster of conserved residues on NTD is buried by the RBD on the spike protein (Figure 5A), but is solvent exposed when NTD is presented alone (Figure 5B). In contrast, the solvent exposed surface of NTD on the spike is much less conserved. Together with our observations above, it is possible that, when immunization is performed using NTD, a reasonable percentage of antibodies are elicited to the conserved surface of NTD that is buried when presented on the spike. Besides, NTD is highly N-glycosylated. It is possible that the N-glycoforms are different between when NTD is expressed alone and when presented on the spike. Such differences may also contribute to the disparity in antigenicity. Therefore, our structural analysis offer an explanation of 1) why NTD immunization elicits antibodies that cross-react with heterologous NTD but not heterologous spike protein (Figure 4A and C), and 2) why immunization with NTD but not live virus, which carries the full spike protein, elicit antibodies that cross-react with heterologous NTD (Figure 2B).

**Figure 5.**
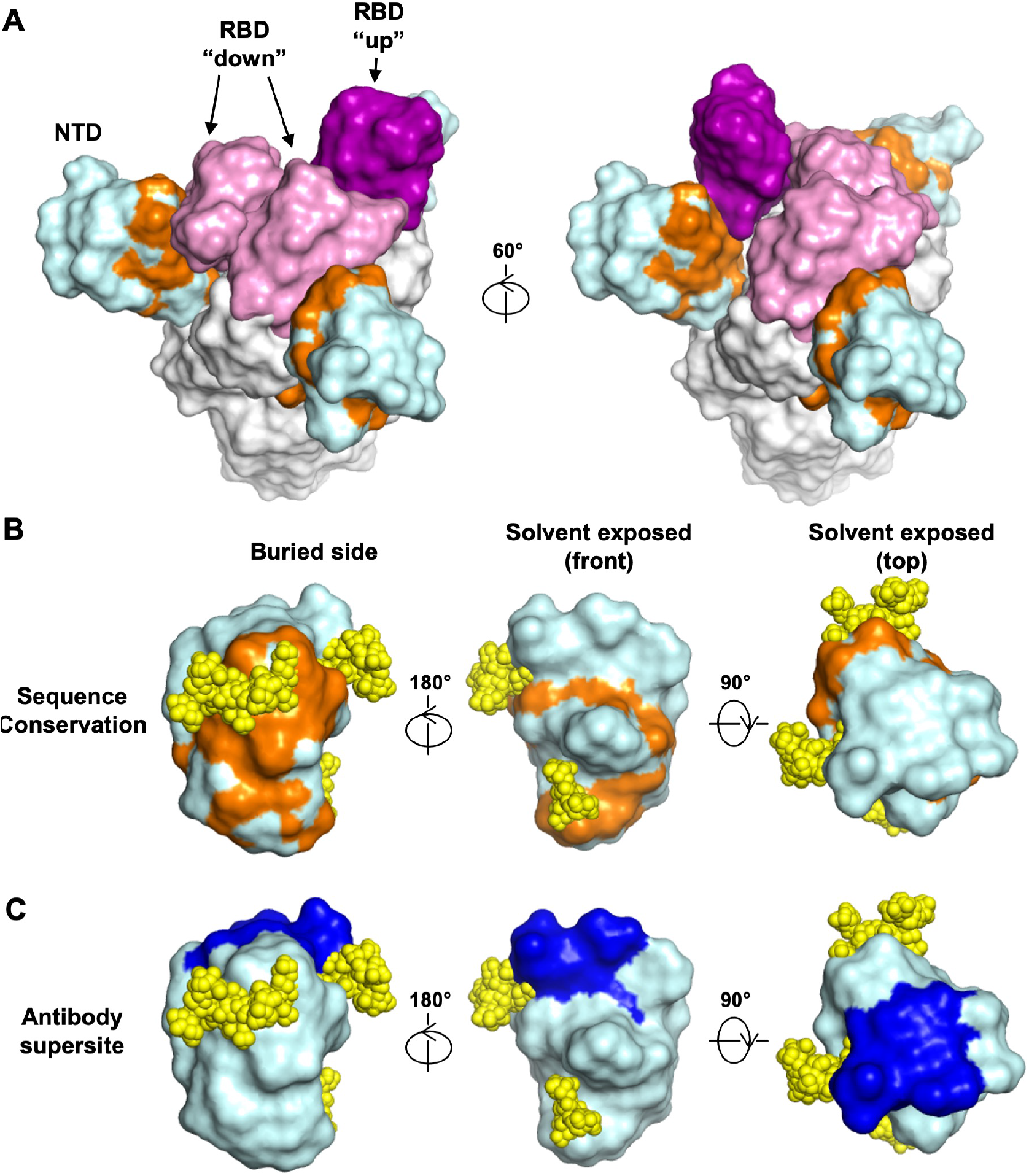
Conservation of NTD protein surface residues between SARS-CoV-2 and SARS-CoV. **(A-B)** Surface residues of NTD (cyan) that are conserved between SARS-CoV-2 and SARS-CoV are highlighted in orange on **(A)** the spike protein where two RBD are in the down conformation (pink) and one RBD is in the up conformation (purple), and on **(B)** NTD alone. **(C)** NTD antibody supersites (McCallum et al., 2021) highlighted in blue. Oligomannoses (yellow) were modeled by GlyProt (Bohne-Lang and von der Lieth, 2005).

## Discussion

Identification of neutralizing antibodies and their targets on SARS-CoV-2 have been a major research area due to the importance for vaccine development. Over the past year, studies have shown that both RBD-specific and NTD-specific antibodies can confer potent neutralizing activity (Barnes et al., 2020; Cerutti et al., 2021; Chi et al., 2020; Ju et al., 2020; Liu et al., 2020a; Liu et al., 2020b; McCallum et al., 2021; Pinto et al., 2020; Seydoux et al., 2020; Suryadevara et al., 2021). However, while SARS-CoV-2 RBD protein can be effective in eliciting neutralizing antibodies (Dai et al., 2020; Tai et al., 2020; Zang et al., 2020), our studies shows that NTD protein is a poor immunogen for eliciting neutralizing antibodies since its antigenicity is altered when expressed alone, where responses may be elicited to epitopes on the NTD that are inaccessible in the spike protein.

Nevertheless, NTD protein can be a useful tool for serology testing. After SARS-CoV-2 infection, both RBD and NTD binding antibodies can be induced in the patient plasma samples after day 14 of symptom onset, suggesting both proteins are suitable for serology testing. In fact, the SARS-CoV-2 RBD protein has been using for serological diagnosis (Perera et al., 2020; Premkumar et al., 2020). However, RBD-specific antibodies can be cross-reactive among SARS-CoV, SARS-CoV-2 and other *Sarbecoviruses* and may result in false-positives (Cui et al., 2019; Lv et al., 2020a; Rappazzo et al., 2021). Moreover, several cross-reactive epitopes against RBD also have been identified between SARS-CoV and SARS-CoV-2 (Liu et al., 2020a; Pinto et al., 2020; Yuan et al., 2020). In contrast, our results show that the cross-reactivity of NTD-specific antibodies to SARS-CoV is much lower that RBD-specific antibodies, indicating that NTD protein could minimize false positives and be an alternative for SARS-CoV-2 serology testing.

One interesting finding in our study is that some SARS-CoV-2 infected patients showed elevation of serological antibody responses against the spike proteins of another two human coronaviruses, HKU1 and OC43. Immunological imprinting in SARS-CoV-2 infected patients due to previous seasonal human coronavirus infection has also been reported (Anderson et al., 2020; Aydillo et al., 2020). Consistently, two conserved cryptic epitopes located in the S2 domain have recently discovered that enable cross-neutralization among five human-infecting beta-coronaviruses, including SARS-CoV, SARS-CoV-2, MERS and OC43 (Huang et al., 2021; Sauer et al., 2021; Song et al., 2020). These observations open up the possibility to develop a more universal vaccine for beta-coronaviruses.

## Method detail

### Virus and Cell cultures

Vero and Vero E6 cells were maintained in DMEM medium supplemented with 10% fetal bovine serum (FBS), and 100 U mL^−1^ of Penicillin-Streptomycin. Sf9 cells (*Spodoptera frugiperda* ovarian cells, female) and High Five cells (*Trichoplusia ni* ovarian cells, female) were maintained in HyClone insect cell culture medium.

Patient-derived SARS-CoV-2 (BetaCoV/Hong Kong/VM20001061/2020 [KH1]) and SARS-CoV (strain HK39849, SCoV) were passaged in Vero-E6 or Vero cells. The virus stock was aliquoted and titrated to determine tissue culture infection dose 50% (TCID_50_). The neutralization experiments were carried out in a Bio-safety level 3 (BSL-3) facility at the School of Public Health, LKS Faculty of Medicine, The University of Hong Kong.

### Collection of specimens

Specimens of heparinized blood were collected from the RT-PCR-confirmed COVID-19 patients at the Infectious Disease Centre of the Princess Margaret Hospital, Hong Kong. All study procedures were performed after informed consent. Plasma from healthy blood donors were collected from the Hong Kong Red Cross before the first COVID-19 case reported on 1^st^ December 2019 (March 2018 to November 2019). The study was approved by the institutional review board of the Hong Kong West Cluster of the Hospital Authority of Hong Kong (approval number: UW20-169). Day 1 of clinical onset was defined as the first day of the appearance of clinical symptoms. The blood samples were centrifuged at 3000 xg for 10 minutes at room temperature for plasma collection. All plasma was kept in −80°C until used.

### Mouse immunization

6-10 weeks BALB/c mice were immunized with two rounds either 15ug NTD protein or 10^5^ TCID_50_ live viruses together with 50 μL Addavax, via intraperitoneal (i.p.) route. The boost dose was given to the mice 21 days after the first priming. The plasma samples were collected using heparin tubes on day 14 after the second round of immunization. The experiments were conducted in The University of Hong Kong Biosafety Level 3 (BSL3) facility. The study protocol was carried out in strict accordance with the recommendations and was approved by the Committee on the Use of Live Animals in Teaching and Research of the University of Hong Kong (CULATR 5422-20).

### Protein expression and purification

The ectodomain (residues 14-1213) with R682G/R683G/R685G/K986P/V987P mutations, receptor-binding domain (RBD, residues 319–541) and N-terminal domain (NTD, residues 14 to 305) of the SARS-CoV-2 spike protein (GenBank: QHD43416.1), as well as the ectodomain (residues 14-1195) with K968P/V969P mutations, RBD (residues 306-527) and NTD (residues 14-292) of the SARS-CoV spike protein (GenBank: ABF65836.1) were cloned into a customized pFastBac vector (Lv et al., 2020b; Wec et al., 2020). The RBD and NTD constructs were fused with an N-terminal gp67 signal peptide and a C-terminal His6 tag. Recombinant bacmid DNA was generated using the Bac-to-Bac system (Life Technologies, Thermo Fisher Scientific). Baculovirus was generated by transfecting purified bacmid DNA into Sf9 cells using FuGENE HD (Promega, Madison, US) and subsequently used to infect suspension cultures of High Five cells (Life Technologies) at a multiplicity of infection (MOI) of 5 to 10. Infected High Five cells were incubated at 28 °C with shaking at 110 rpm for 72 h for protein expression. The supernatant was then concentrated using a Centramate cassette (10 kDa molecular weight cutoff for RBD, Pall Corporation, New York, USA). RBD and NTD proteins were purified by Ni-NTA Superflow (Qiagen, Hilden, Germany), followed by size exclusion chromatography and buffer exchange to phosphate-buffered saline (PBS). The spike proteins of 229E, HKU1, NL63 and OC43 were purchased from Sino Biological (China).

### ELISA

A 96-well enzyme-linked immunosorbent assay (ELISA) plate (Nunc MaxiSorp, Thermo Fisher Scientific) was first coated overnight with 100 ng per well of purified recombinant protein in PBS buffer. The plates were then blocked with 100 μl of Chonblock blocking/sample dilution ELISA buffer (Chondrex Inc, Redmon, US) and incubated at room temperature for 1 h. Each human plasma sample was diluted to 1:100 in Chonblock blocking/sample dilution ELISA buffer. Each sample was then added into the ELISA plates for a two-hour incubation at 37°C. After extensive washing with PBS containing 0.1% Tween 20, each well in the plate was further incubated with the anti-human IgG secondary antibody (1:5000, Thermo Fisher Scientific) for 1 hour at 37°C. The ELISA plates were then washed five times with PBS containing 0.1% Tween 20. Subsequently, 100 μL of HRP substrate (Ncm TMB One; New Cell and Molecular Biotech Co. Ltd, Suzhou, China) was added into each well. After 15 min of incubation, the reaction was stopped by adding 50 μL of 2LJM H_2_SO_4_ solution and analyzed on a Sunrise (Tecan, Männedorf, Switzerland) absorbance microplate reader at 450 nm wavelength (Perera et al., 2020).

### Plaque reduction neutralization test (PRNT)

Plasma samples were two-fold diluted starting from a 1:10 dilution and mixed with equal volumes of around 120 plaque-forming units (pfu) of SARS-CoV-2 or SARS-CoV as determined by Vero E6 and Vero cells respectively. After 1-hour incubation at 37°C, the plasma-virus mixture was added onto cell monolayers seated in a 24-well cell culture plate and incubated for 1 hour at 37°C with 5% CO_2_. The plasma-virus mixtures were then discarded and infected cells were immediately covered with 1% agarose gel in DMEM medium. After incubation for 3 days at 37°C with 5% CO_2_, the plates were formalin fixed and stained by 0.5% crystal violet solution. Neutralization titers were determined by the highest plasma dilution that resulted in >90% reduction in the number of pfus. The test was performed in a BSL3 facility at the University of Hong Kong (Perera et al., 2020).

### Statistical analysis

We defined a sample as ELISA antibody positive if the OD value was 3 standard deviations above the mean of the negative controls. Significance between two groups were determined by Mann-Whitney test with p-values lower than 0.05. Correlation between plasma samples were assessed using Pearson correlation coefficients. Two-tailed paired t-tests were performed in Figure 3.

## Acknowledgements

This work was supported by Calmette and Yersin scholarship from the Pasteur International Network Association (H.L.), Bill and Melinda Gates Foundation OPP1170236 and INV-004923 (I.A.W.), startup funds from the University of Illinois at Urbana-Champaign (N.C.W.), the US National Institutes of Health (contract no. HHSN272201400006C) (J.S.M.P), National Natural Science Foundation of China (NSFC)/Research Grants Council (RGC) Joint Research Scheme (N_HKU737/18) (C.K.P.M. and J.S.M.P), the Research Grants Council of the Hong Kong Special Administrative Region, China (Project no. T11-712/19-N) (J.S.M.P) and Guangdong Province International Scientific and Technological Cooperation Projects (grant number 2020A0505100063). We acknowledge the support of the clinicians who facilitated this study, including Drs Wai Shing Leung, Jacky Man Chun Chan, Thomas Shiu Hong Chik, John Yu Hong Chan, Daphne Pui-Lin Lau, and Ying Man Ho, and the dedicated clinical team at Infectious Diseases Centre, Princess Margaret Hospital, Hospital Authority of Hong Kong and the patients who kindly consented to participate in this investigation.

## Author contributions

H.L., N.C.W., J.S.M.P. and C.K.P.M. conceived the research idea, planned the study, obtained research funding, analysed the data and wrote the manuscript. M.Y., H.L., I.A.W. and N.C.W. expressed and purified the proteins. H.L., R.T.Y.S., Y.W., G.K.Y., Q.T, Y.L., W.L., J.W. and W.W.N performed the experiments. O.T.Y.T organized patient recruitment, data collection and sampling.

## Competing Interests

The authors declare no competing interests.

---

**Figure S1.**
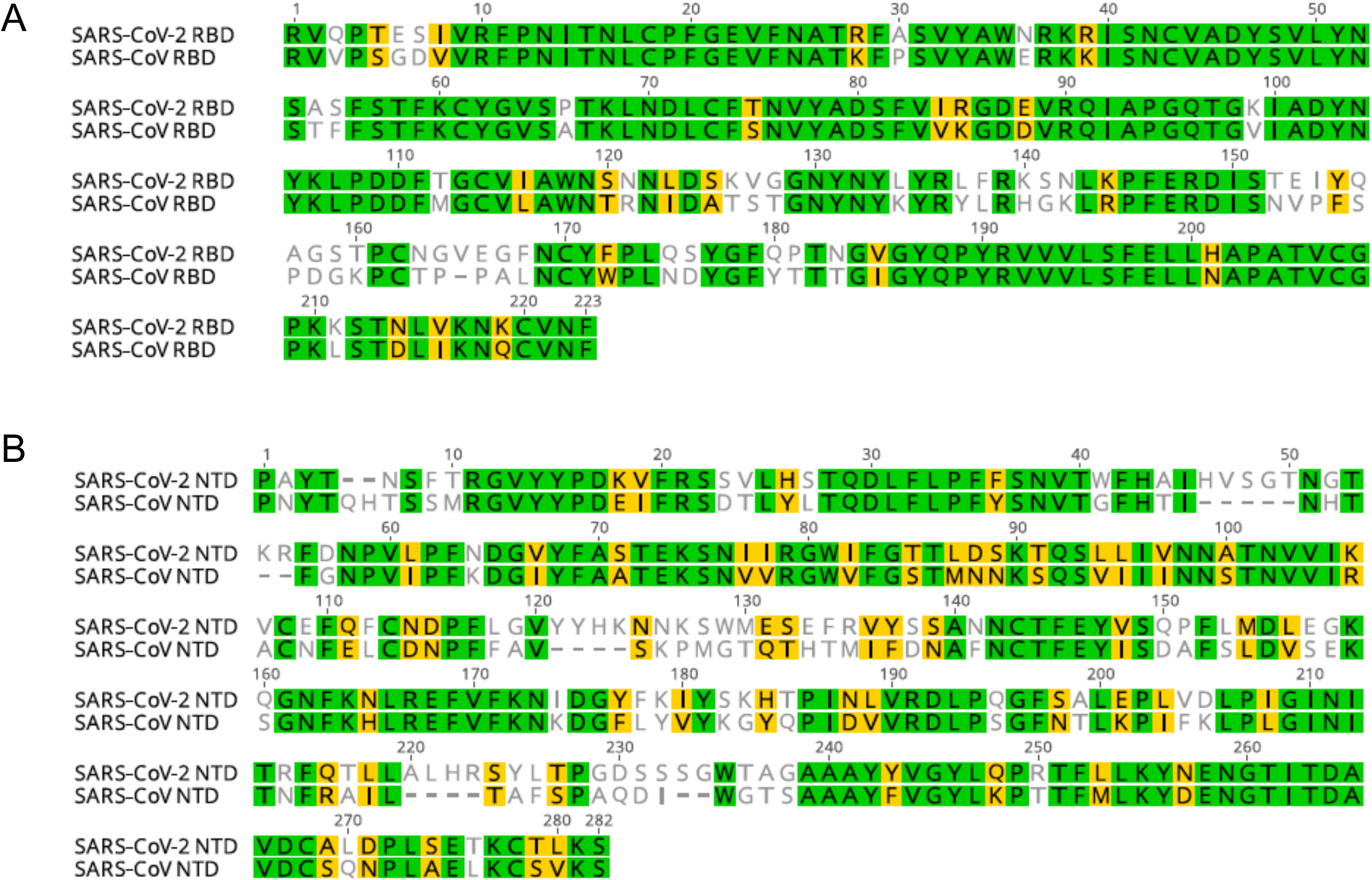
Sequence alignment for SARS-CoV-2 and SARS-CoV RBD and NTD protein. **(A-B)** The RBD and NTD domain of SARS-CoV (NCBI Reference Sequence: NC_004718.3) and SARS-CoV-2 (NCBI Reference Sequence: NC_045512.2) were aligned by MUSCLE (https://www.ebi.ac.uk/Tools/msa/muscle/). Residues highlighted with green and yellow represent identical and similar residues respectively. The percentage identity and similarity are calculated with a Blosum 62 score matrix using Geneious Prime.

**Figure S2.**
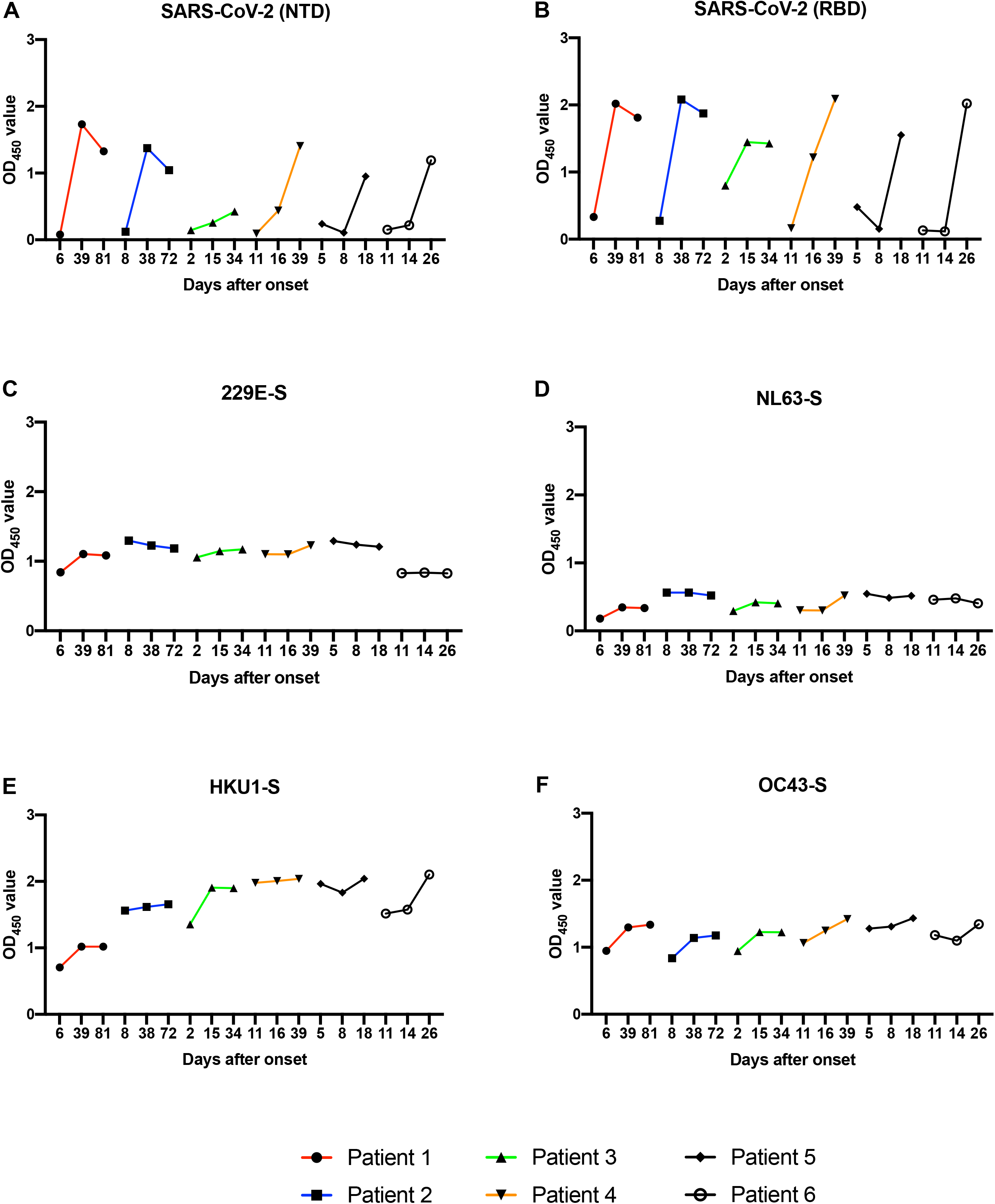
Longitudinal serological analysis to human coronaviruses. **(A-F)** Binding of longitudinal plasma samples against SARS-CoV-2 NTD (A), SARS-CoV-2 RBD **(B)**, 229E-Spike **(C)**, NL63-Spike **(D)**, HKU1-Spike **(E)** and OC43-Spike protein **(F)** from six COVID-2019 patients were measured by ELISA.

## Notes

### Competing Interest Statement

The authors have declared no competing interest.

